# Tau-mediated axonal degeneration is prevented by activation of the Wld^S^ pathway

**DOI:** 10.1101/2020.12.06.408583

**Authors:** Katy Stubbs, Megan Sealey, Miguel Ramirez Moreno, V Hugh Perry, Tracey A Newman, Amritpal Mudher

## Abstract

Tauopathy is characterised by neuronal dysfunction and degeneration occurring as a result of changes to the microtubule associated protein tau. The neuronal changes evident in Tauopathy bear striking morphological resemblance to those reported in models of Wallerian degeneration. The mechanisms underpinning Wallerian degeneration are not fully understood although it can be delayed by the expression of the slow Wallerian degeneration (Wld^S^) protein, which has also been demonstrated to delay axonal degeneration in some models of neurodegenerative disease. Given the morphological similarities between tauopathy and Wallerian degeneration, this study investigated whether tau-mediated phenotypes can be modulated by expression of Wld^S^. In a *Drosophila* model of tauopathy in which expression of human Tau protein (hTau^0N3R^) leads to progressive age-dependent phenotypes, activation of the pathway downstream of Wld^S^ completely suppressed tau-mediated degeneration. This protective effect was evident even if the pathway downstream of Wld^S^ was activated several weeks after hTau-mediated degeneration had become established. In contrast, Wld^S^ expression without activation of the downstream protective pathway did not rescue tau-mediated degeneration in adults or improve tau-mediated neuronal dysfunction including deficits in axonal transport, synaptic alterations and locomotor behaviour in hTau^0N3R^ –expressing larvae. This collectively implies that the pathway mediating the protective effect of Wld^S^ intersects with the mechanism(s) of degeneration initiated by hTau and can effectively halt tau-mediated degeneration at both early and late stages. Understanding the mechanisms underpinning this protection could identify much-needed disease-modifying targets for tauopathies.

## Introduction

Tau pathology is observed in numerous neurodegenerative diseases, including Alzheimer’s disease, Parkinson’s disease (PD), motor neuron disease (MND) and a variety of other tauopathies such as fronto-temporal dementia, Pick’s Disease, progressive supra-nuclear palsy and others. The axon is susceptible to tau pathology in these neurodegenerative diseases, with evidence of white matter changes indicative of axonal degeneration in tauopathies such as AD (1-3). Studies in animal models have demonstrated that axonal dysfunction in tauopathy is typified by disrupted axonal transport (4, 5) (6), due to tau hyperphosphorylation resulting in reduced cytoskeletal integrity (7). Axonal swellings and loss of white matter, hallmarks of axonal degeneration have been observed in P301L-tau mice, a model of familial fronto-temporal dementia (8) (9) (10).

Wallerian degeneration describes the sequential degeneration of axons following axonal injury which begins with breakdown of the cytoskeleton and ends with the fragmentation and loss of the separated distal axon (11). Wallerian degeneration and axonal degeneration in neurodegenerative disease share similarities including cytoskeletal breakdown (7) (12), disrupted axonal transport (13) (14), alterations to mitochondrial morphology (15) (16), and in the central nervous system (CNS), axonal swellings (12) (17). These similarities suggest that the mechanisms overlap and the term Wallerian-like may be used to describe degeneration that is not due to an acute injury.

The discovery of the slow Wallerian degeneration (Wld^S^) protective mutation, which robustly delays Wallerian degeneration (18), identified a molecular pathway controlling axonal degeneration after injury [reviewed in (19) (20) (21) (22)]. Wld^S^ has been studied in experimental models to elucidate the molecular pathway and explore whether it underpins the Wallerian-like degeneration observed in a variety of neurodegenerative conditions. This work has identified delayed degeneration in models of disease including: Multiple sclerosis (23), Parkinson’s disease (24) (25), Charcot-Marie-Tooth disease type 1A (26) and 1B (27) and toxic neuropathy (28).

Considering that the axon is a site of tau-mediated dysfunction and degeneration, the aim of the present study was to investigate whether the axonal protection mediated by Wld^S^ was able to rescue tau-mediated axonal dysfunction and degeneration. *Drosophila melanogaster* has been used in the study of Wallerian degeneration and Wld^S^ (29) (30) (31) (32) (33) (34), and *Drosophila* models of tauopathy are similarly well-established (5) (35) (36) . Furthermore, several studies are beginning to implicate components of Wld^S^ (such as nicotinamide mononucleotide adenylyl transferase – NMNAT) in tau-mediated aggregation and degeneration in both rodent and *Drosophila* models (37) (38) (39). To investigate whether Wallerian-like degeneration in tauopathy is Wld^S^-sensitive we studied the structural and functional effects of co-expression of human tau (hTau^0N3R^) and Wld^S^ in *Drosophila*.

Our findings demonstrate that co-expression of Wld^S^ does not confer protection against hTau mediated dysfunction or degeneration. However, in stark contrast, activation of the pathway downstream of Wld^S^ results in profound protection, both preventing and arresting degeneration even in neurons already affected by tau-induced pathology.

## Materials and methods

### Fly stocks

*Drosophila* were raised and maintained on standard Bloomington media at 23°C with a 12/12 h light/dark cycle. UAS-htau^0N3R^, elav-GAL4, D42-GAL4 and Oregon R flies were obtained from the Bloomington Drosophila Stock Centre (Indiana, IN, USA). The UAS-Wld^S^ and UAS-mCD8::GFP, Or47b-GAL4/Cyo lines were obtained from Professor Liqun Luo (Stanford University, CA, USA (40)). The D42-GAL4, UAS-NPY::GFP line was generated previously (5), with UAS-NPY::GFP provided by Dr Ian Robinson (Plymouth University, UK). A homozygous htau^0N3R^;Wld^S^ line was generated for the current study by crossing UAS-htau^0N3R^ with UAS-Wld^S^ lines.

### Axonal transport analysis

Transgenes were expressed using D42-GAL4, UAS-NPY::GFP, and third instar wandering larvae were selected for analysis. Larvae were anaesthetised using diethyl ether vapour (Thermo Fisher Scientific) and mounted in 1% agarose (Sigma-Aldrich) on glass slides, with the ventral surface facing the coverslip. Peripheral nerves were imaged using an Axioplan2 MOT upright fluorescence microscope (Zeiss) equipped with Micro Max CCD (Princeton Instruments) using MetaMorph acquisition software (Molecular Devices). Images were thresholded and the area covered by aggregates measured using Metamorph software.

### Larval neuromuscular junction (NMJ) analysis

Transgenes were expressed using D42-GAL4, and third instar wandering larvae dissected, with internal organs removed and the skin pinned out and fixed in 4% formaldehyde (Sigma-Aldrich) for 90 mins at room temperature. Larval skins were then washed in 0.1% Triton X (Sigma-Aldrich) in phosphate-buffered saline (PBS-Tx; Thermo Fisher Scientific) prior to blocking in 5% goat serum, 3% horse serum and 2% bovine serum albumin (BSA; Sigma-Aldrich) in 0.1% PBS-Tx. Skins were incubated with goat anti-horseradish peroxidase (1:1000; ICN/Cappel), conjugated to fluorescein isothiocyanate. Skins were washed in 0.1% PBS-Tx and put through an ascending glycerol series (50, 70, 90 and 100%) before being mounted in Vectashield (Vector Laboratories) and imaged. NMJ’s on muscle 4 from segments A3-5 were imaged using a Leica SP2 scanning confocal microscope using the 488 argon laser. Maximum projections of Z stacks were generated for morphometric analysis; bouton size and interbouton axon width were measured using ImageJ with the assessor blinded to the sample number.

### Larval locomotion

Larval behaviour was assessed as previously described (41). In brief, D42-GAL4 driven third instar wandering larvae were each placed in the centre of 0.3% Alsian Blue (Sigma-Aldrich), 1% agarose (Sigma-Aldrich) plates, and videos of larval behaviour recorded. Videos of larval locomotion were analysed using Ethovision 3.0 (Noldus) tracking software.

### Immunohistochemical analysis of axonal degeneration

Adults were collected 0-2 days after eclosion from UAS-mCD8::GFP, Or47b-GAL4 driven crosses and aged to the relevant time point. Flies were anaesthetised with CO2, heads were ligated and the brains dissected and placed in 4% formaldehyde and fixed at room temperature for 45 minutes. Following fixation, brains were washed in 0.1% PBS-Tx before either mounting in Vectashield or proceeding for staining. Those to be stained were blocked in 5% goat serum, 3% horse serum, 2% BSA in 0.1% PBS-Tx and stained with rabbit anti-human tau antibodies (1:1000; Dako) or mouse anti-phospho tau PHF-1 (1:1000 – thermofisher), washed and incubated in goat anti-rabbit or anti-mouse Alexa Fluor 563 (1:1000; Invitrogen; Thermo Fisher Scientific). Brains were washed and mounted in Vectashield prior to imaging on an Axioplan2 MOT upright Epifluorescence microscope (Zeiss) equipped with a QImaging Retiga 3000 CCD Camera (Photometrics) and images were acquired using Metamorph software (Molecular Devices). Images were quantified in ImageJ with the assessor blinded to genotype and timepoint. For axonal swellings, images were thresholded and the coverage of swellings measured.

### Axon injury to activate the pathway downstream of Wld^S^ (referred to as Wld^S^ pathway-activation)

The third antennal segment was removed from flies, 1 or 3 weeks after eclosion from Or47b-GAL4 driven crosses, under CO2 anaesthesia using Dumont #5 forceps. This induced an axonal injury in olfactory receptor neurons (ORNs), whose cell bodies are located in the third antennal segment. At the relevant time points, brains were dissected as described above. Degeneration was quantified by previously described methods (42). Briefly, with the assessor blind to genotype and time point, the presence of the commissural axons was recorded (Y/N) and the percentage of brains of each genotype at each time point with intact axons was calculated. The intensity of GFP signal within glomeruli was measured using ImageJ and the background intensity was subtracted.

### Statistics

Statistical analysis was conducted using GraphPad Prism, version 6.0 (GraphPad Software, Inc.), using analysis of variance and the Bonferroni correction for the comparison of groups. The Mantel-Cox test was used for survival analysis, with the Bonferroni correction used for the comparison of multiple groups. Values are presented as the mean ± standard error. P<0.05 was considered to indicate a statistically significant difference.

## Results

### Activation of the pathway downstream of Wld^S^ protects against hTau^0N3R^-induced degeneration

Though previous studies of injury models indicate that the presence of Wld^S^ within the axon is crucial for its protection (33-35), the findings from chronic models of disease do not show any consistent or significant Wld^S^-mediated protection despite clear evidence of Wallerian-like degeneration in these models (31) (32) (34). One explanation for this lack of rescue could be that the pathway that Wld^S^ is acting in is not “activated” in these models of chronic degeneration, raising the possibility that the protein may require some form of injury to unmask its protective effect. Indeed, in all cases where Wld^S^ has been reported to rescue axonal degeneration, the neurons are injured by default as part of the experimental paradigm (22) (26) (35) (36).

To explore whether injury-induced “activation” was required for the Wld^S^ pathway to protect against tau-mediated degeneration, hTau^0N3R^;Wld^S^ axons of the olfactory receptor neurons (ORNs) expressing membrane bound GFP were injured (axotomised) by removal of the third antennal segments as previously described (42). Adult brains were analysed after ecclosion at hourly (h) or weekly (w) time points post-axotomy induced Wld^S^ pathway activation (referred to as “pa” from here on). This revealed that control and tau expressing axons had degenerated at 2w after eclosion/1wpa. In contrast hTau^0N3R^;Wld^S^ expressing axons were intact at this time point (data not shown) confirming that the axotomy paradigm “activated” the pathway downstream of Wld^S^. To ascertain the extent to which activation of the Wld^S^ pathway protected against tau-mediated degeneration, the prominent degenerative features of tau-expressing axons were quantified. Axonal swellings, which are characteristic of tau-mediated axonal degeneration, were present in naïve hTau^0N3R^;Wld^S^ axons (arrowheads Fig 1ai). In contrast these were not found in hTau-animals of the same genotype after activation of the Wld^S^ pathway (Fig. 1aii). The progressive accumulation of axonal swellings is evident in hTau expressing animals within 2 weeks after eclosion and trebles by week 5. In contrast swellings were not seen at any time point in hTau^0N3R^;Wld^S^ expressing animals where the activation of the Wld^S^ pathway was elicited through axotomy (Fig. 1b). This illustrates that once activated, the Wld^S^ pathway protects against the initiation and development of tau-mediated axonal degeneration.

**Fig 1.**
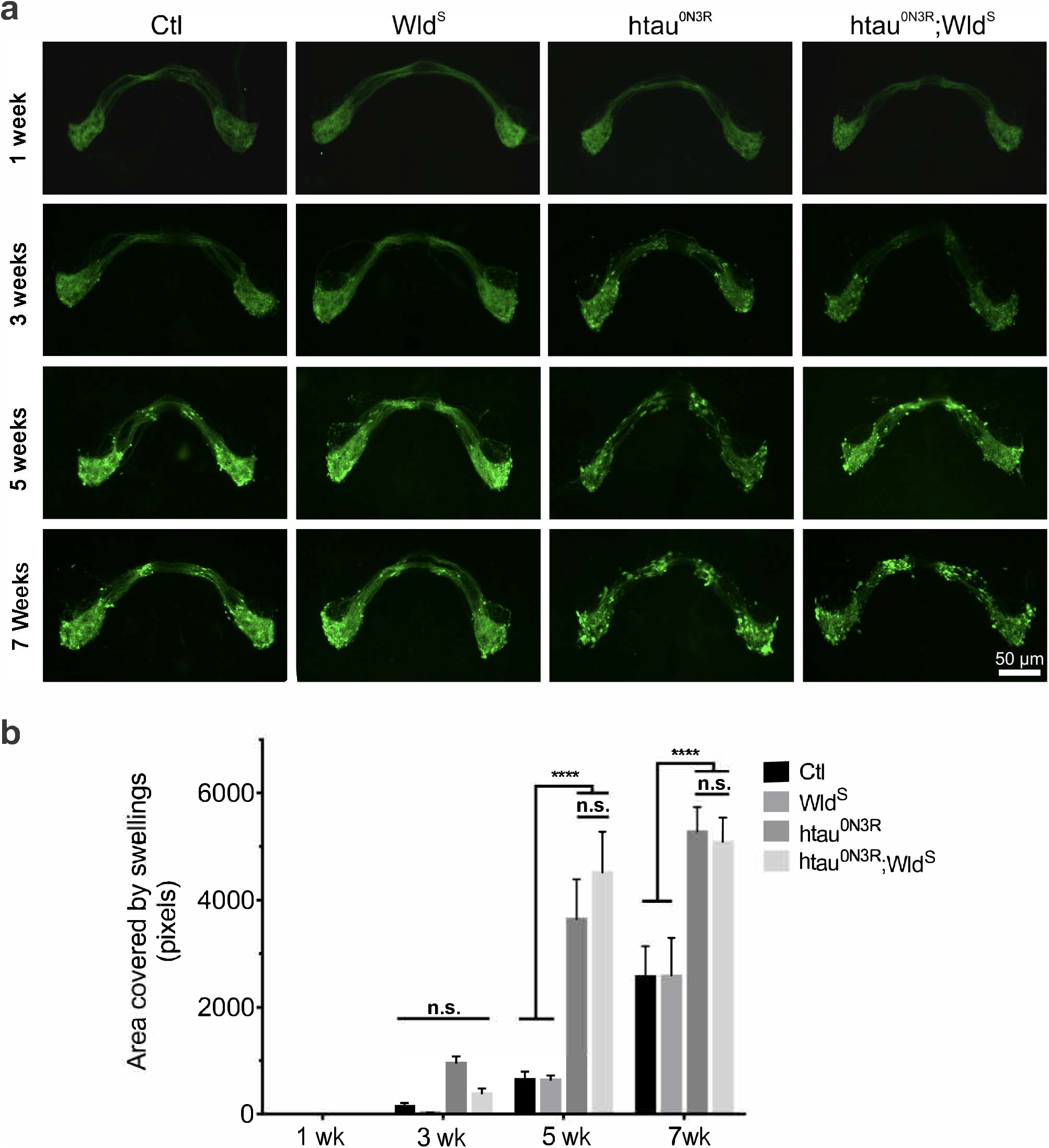
Tau-mediated axonal swellings are not evident in hTau^0N3R^;Wld^S^ axons in which Wld^S^ pathway is activated. a) Axonal swellings (arrowheads) in naïve htau^0N3R^;Wld^S^ axons where Wld^S^ pathway has not been activated. b) These swellings do not appear in hTau^0N3R^;Wld^S^ axons where Wld^S^ has been activated, at any time point post-activation (pa). b) Quantification of coverage of axonal swellings. Values are presented as the mean ± SEM. n=6-10. P<0.0001.

To study a more disease-relevant situation we investigated whether activation of the Wld^S^ pathway protected against already established tau-mediated axonal degeneration. The Wld^S^ pathway was activated at 3w after eclosion, a time at which axonal swellings are already established in hTau^0N3R^-expressing axons. In hTau^0N3R^;Wld^S^ axons where Wld^S^ pathway was not activated, a progressive increase in axonal swellings is evident with time, such that axonal swellings at 4w after eclosion are 4-fold greater than those seen at 3w after eclosion, with this increasing further by 6w after eclosion (P<0.001; Fig 2a). At these later time points the swellings in the naïve hTau^0N3R^;Wld^S^ axons are significantly greater than those seen in the naïve Wld^S^ axons which serve as the controls. In contrast activation of the Wld^S^ pathway halts the development of axonal swelling. Olfactory receptor neurons in flies expressing hTau;Wld^S^ showed the anticipated accumulation of axonal swellings at 3w after eclosion, prior to Wld^S^ pathway activation, but any further accumulation was halted once the Wld^S^ pathway was activated (Fig. 2b) with no progression in pathology seen after this time. Once the pathway downstream of Wld^S^ was activated, the axonal swellings in hTau^0N3R^;Wld^S^ animals were not significantly different to those seen in controls at any time point. This indicates that in addition to preventing the emergence of tau-mediated axonal degeneration (as shown in Fig 1), activation of the Wld^S^ pathway can also halt the progression of tau-mediated axonal degeneration once it has begun.

**Fig 2.**
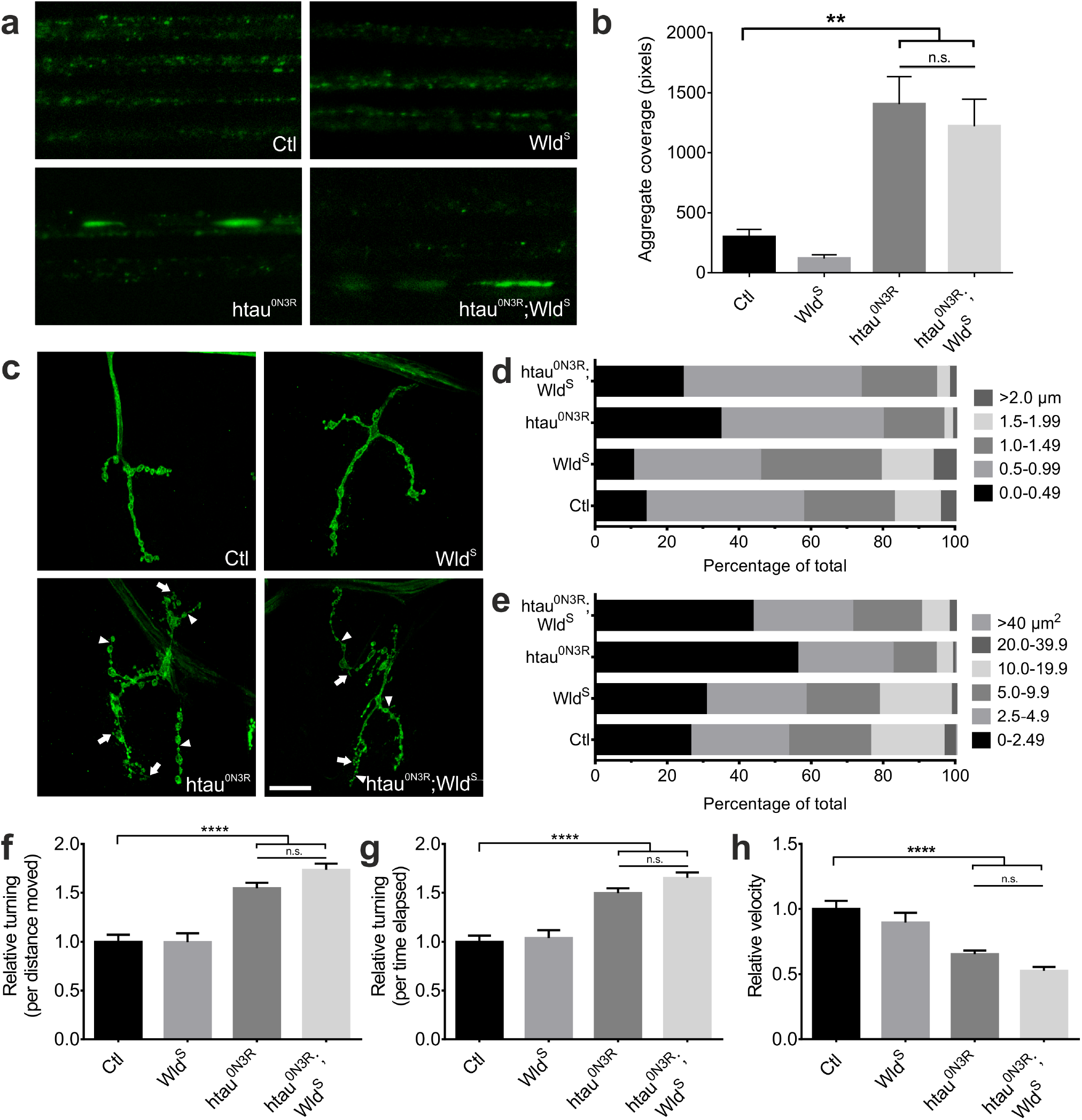
Tau-mediated axonal swellings are halted from progressing upon activation of the Wld^S^ pathway. a) Quantification of axonal swellings reveals that in naïve hTau^0N3R^;Wld^S^ axons, where Wld^S^ pathway has not been activated, the level of swellings increases significantly at 4 wks and 6 wks (n=7-8). b) In hTau^0N3R^;Wld^S^ axons where Wld^S^ has been activated the area covered by axonal swellings does not increase significantly over time post Wld^S^ pathway activation (pa) (n=8-16). Values are presented as the mean ± SEM. ***P<0.001, ****P<0.0001.

### Expression of Wld^S^ without “activation” of the downstream pathway is insufficient to protect against hTau^0N3R^-induced phenotypes

The results indicate that activation of the pathway downstream of Wld^S^ potently suppresses hTau-mediated degeneration. This supports our hypothesis that the lack of protection through co-expression of Wld^S^ in chronic models of degeneration (31) (32) (33) (34) is because the pathway that Wld^S^ acts in is not normally “activated” in otherwise naïve axons. Acute injury to an axon “activates” it unmasking its protective effect. However, as the previous studies were conducted in rodents, we sought to ascertain whether the “protection requires pathway activation” phenomenon that we described in Figs 1 and 2 holds true in our invertebrate model as well.

To prove that expression of Wld^S^ is insufficient for protection against hTau^0N3R^-mediated degeneration and that “activation” of the pathway downstream is required, degeneration in naïve hTau^0N3R^;Wld^S^ flies was studied in the absence of “activation” . ORNs expressing membrane-bound GFP underwent progressive age-related axonal degeneration in all hTau^0N3R^ expressing flies. This was characterised by the appearance of axonal swellings at 2-3 weeks after eclosion, which increased in number and size as the flies aged (Fig. 3a/b hTau^0N3R^ column). Axonal swellings were also evident in controls and Wld^S^ flies, but only at older, 5-7 week time points (Fig. 3a control and Wld^s^ columns). Noticeably, these swellings were also apparent in hTau^0N3R^;Wld^S^ flies (where the Wld^s^ pathway had not been activated - Fig 3a hTau^0N3R^;Wld^s^ column). Quantification confirmed that there was no significant difference in onset, extent or progression of axonal swellings in the hTau^0N3R^;Wld^S^ flies when compared with hTau^0N3R^ alone (Fig. 3a/b).

**Fig 3.**
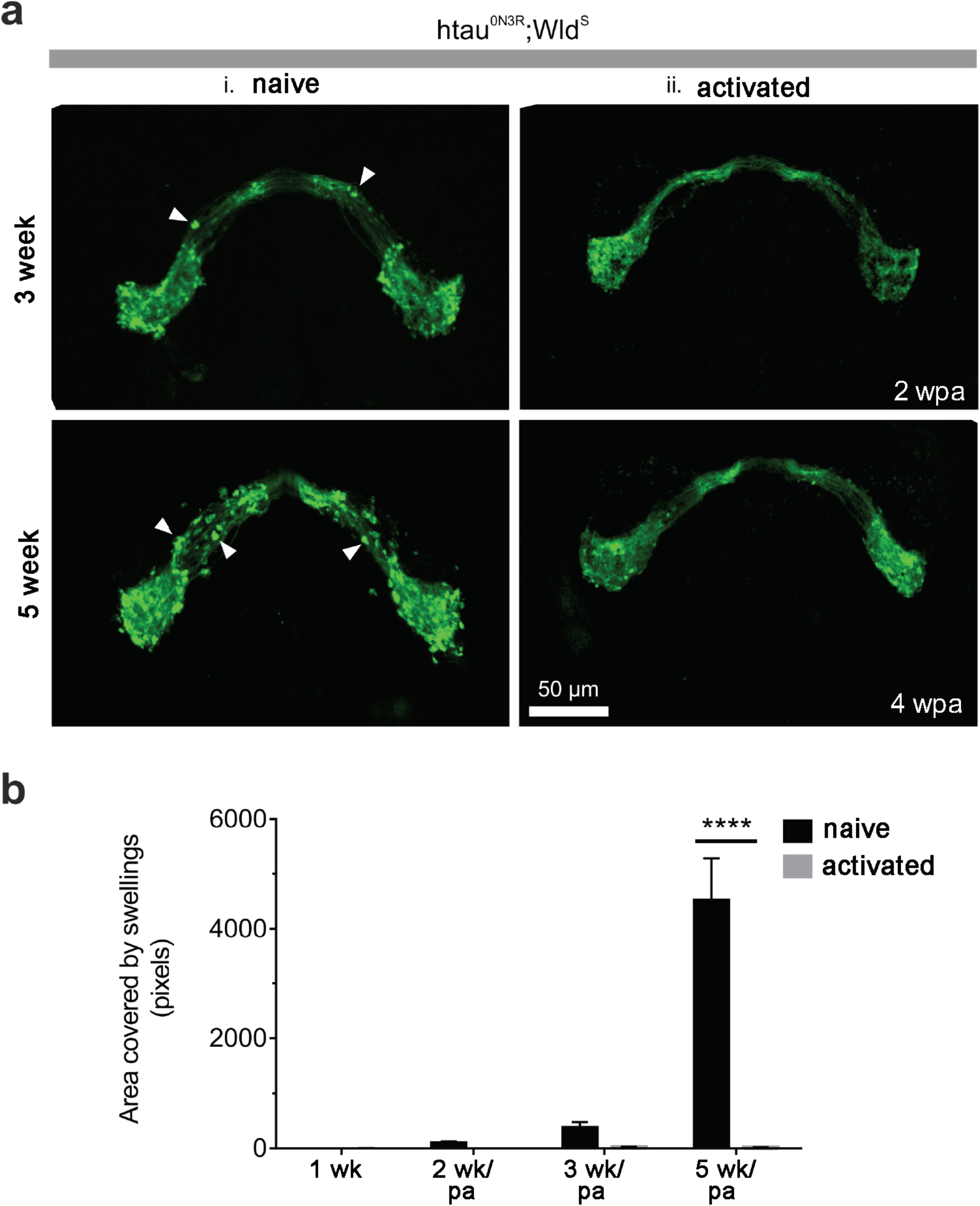
Co-expression of Wld^S^ with hTau^0N3R^ does not delay tau-mediated axonal degeneration. At 1 week after eclosion all genotypes display normal olfactory receptor neuron (ORN) morphology (a). At 3 weeks after ecclosion, axonal swellings are apparent in htau^0N3R^ expressing ORNs with similar morphology observed in hTau^0N3R^;Wld^S^ ORNs. b) Quantification of swelling coverage indicates that co-expression of Wld^S^ does not delay the onset nor slow the progression of tau-mediated axonal degeneration. Values are presented as the mean ± SEM. n=6-11, ****P<0.0001. Wld^S^, slow Wallerian degeneration; hTau^0N3R^, 0N3R human tau isoform; n.s., not significant.

These results suggest that simply co-expressing Wld^S^ does not protect against tau-mediated degeneration. We next sought to investigate whether this is also the case in larvae, where tau-mediated neuronal dysfunction manifests in profound Wallerian-like axonal phenotypes incuding disrupted axonal transport and destabilisation of the cytoskeleton (5) (7). Using a *Drosophila* line expressing GFP-tagged neuropeptide Y in motor neurons, axonal transport was visualised using microscopy in live intact third instar larvae. As reported previously, numerous large vesicular aggregates were found in tau-expressing larvae indicative of axonal transport disruption (Fig. 4a). However, these aggregates were also observed in hTau^0N3R^;Wld^S^ larvae. Quantification of the coverage areas of the aggregates indicated that the aggregates were not significantly reduced in hTau^0N3R^;Wld^S^ larvae compared with hTau^0N3R^ larvae (Fig. 4b).

**Fig 4.**
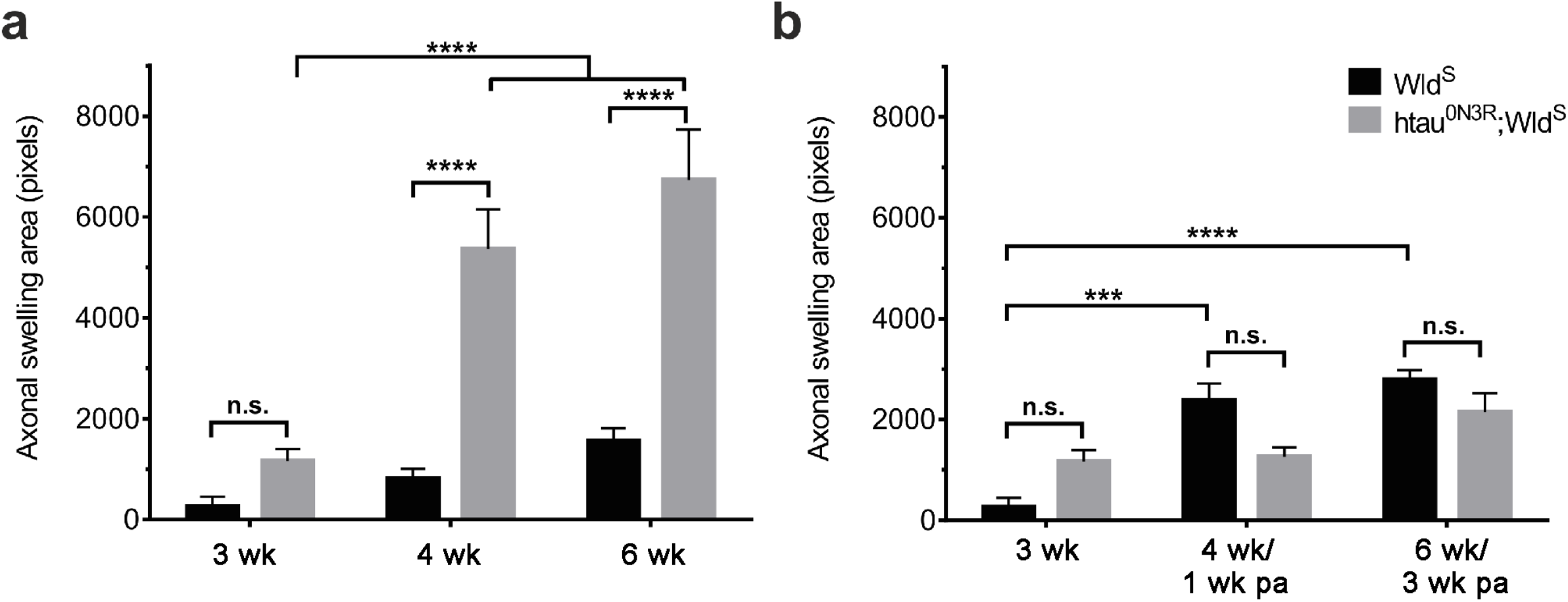
Wld^S^ does not improve tau-mediated axonal dysfunction. a) Expression of hTau^0N3R^ results in the appearance of vesicular aggregates, which are also apparent in hTau^0N3R^;Wld^S^ axons. b) No significant improvement in aggregate coverage was observed in hTau^0N3R^;Wld^S^ axons compared with hTau^0N3R^ axons (n=10 larvae). c) hTau^0N3R^ NMJs display aberrant morphology, typified by thinning of the axon (arrowheads) and microsatellite boutons (arrows), with this also observed in htau^0N3R^;Wld^S^ NMJs (scale bar = 25 µm). Co-expression of Wld^S^ with hTau^0N3R^ did not rescue thinning of the inter-bouton axon (d) or the alterations in bouton size (e) (n=4 larvae, 3-6 NMJs/larva). Analysis of locomotor behaviour indicated that co-expression of hTau^0N3R^;Wld^S^ did not significantly improve the hTau^0N3R^-mediated alterations in f) meander – relative turning/distance travelled, g) angular velocity – relative turning / time elapsed and h) velocity of larval crawling (n>17). Values are presented as the mean ± SEM. **P<0.01; ****P<0.0001. Wld^S^, slow Wallerian degeneration; htau^0N3R^, 0N3R human tau isoform; NMJs, neuromuscular junctions; Ctl, control.

HTau^0N3R^ expression is associated with altered synaptic morphology, characterised by thinning of the inter-bouton axons and the appearance of minisatellite boutons (43). These features were observed in the current study (Fig. 4c) and this phenotype was not improved by co-expression of Wld^S^. No significant difference in the thickness of the inter-bouton axon (Fig. 4d) or the proportions of bouton of each size (Fig. 4e) were seen in the neuromuscular junctions of either hTau^0N3R^;Wld^S^ or hTau^0N3R^ expressing animals .

Disruption of axonal transport and altered synaptic morphology are associated with alterations in locomotor behaviour. Using an open field behavioural assay, the crawling behaviour of third instar larvae was investigated. When placed in the centre of an arena, control larvae quickly move towards the edge of the arena, following a straight path. In contrast, tau expressing larvae move more slowly and take a confused and twisting path, demonstrated by an increase in meander (turning per distance moved; Fig. 4f) and in angular velocity (turning per time elapsed; Fig. 4g) and the reduction in overall velocity (Fig. 4h). However, the co-expression of Wld^S^ with hTau^0N3R^ did not improve locomotor behaviour, with no significant difference between hTau^0N3R^;Wld^S^ and hTau^0N3R^ expressing larvae (Fig. 4f-h).

The adult and larval data collectively shows that without “activation” of the pathway downstream of Wld^S^, the protective effect of Wld^S^ on hTau^0N3R^-mediated dysfunction (in larvae) or degeneration (in adult flies) is not uncovered.

### Activation of the Wld^S^-pathway protects against the effects of hTau^0N3R^ without influencing total or phosphorylated tau levels

The most parsimonious explanation for this curious phenomenon of injured-activated protection against hTau^0N3R^ pathology, may simply be that hTau is lost from injured hTau^0N3R^;Wld^S^ axons and therefore cannot exert it’s detrimental effects to cause axonal degeneration. To investigate this, hTau immunoreactivity was assessed in hTau-expressing animals with and without Wld^S^-pathway activation and both the amount of hTau and its cellular localisation was examined. No significant differences were found in hTau distribution or total hTau expression between these two groups; hTau staining persisted in injured hTau^0N3R^;Wld^S^ axons even 5 weeks after Wld^S^ activation (Fig. 5a) and there was no difference in total Tau levels (Fig 5b). This is remarkable because it implies that despite expression within the axon for 6 weeks, hTau has not caused degeneration in the hTau^0N3R^;Wld^S^ axons once the Wld^S^ pathway is activated. This begs the question as to how activation, of the Wld^S^-pathway protects against the human tau induced degeneration across this length of time.

**Fig 5.**
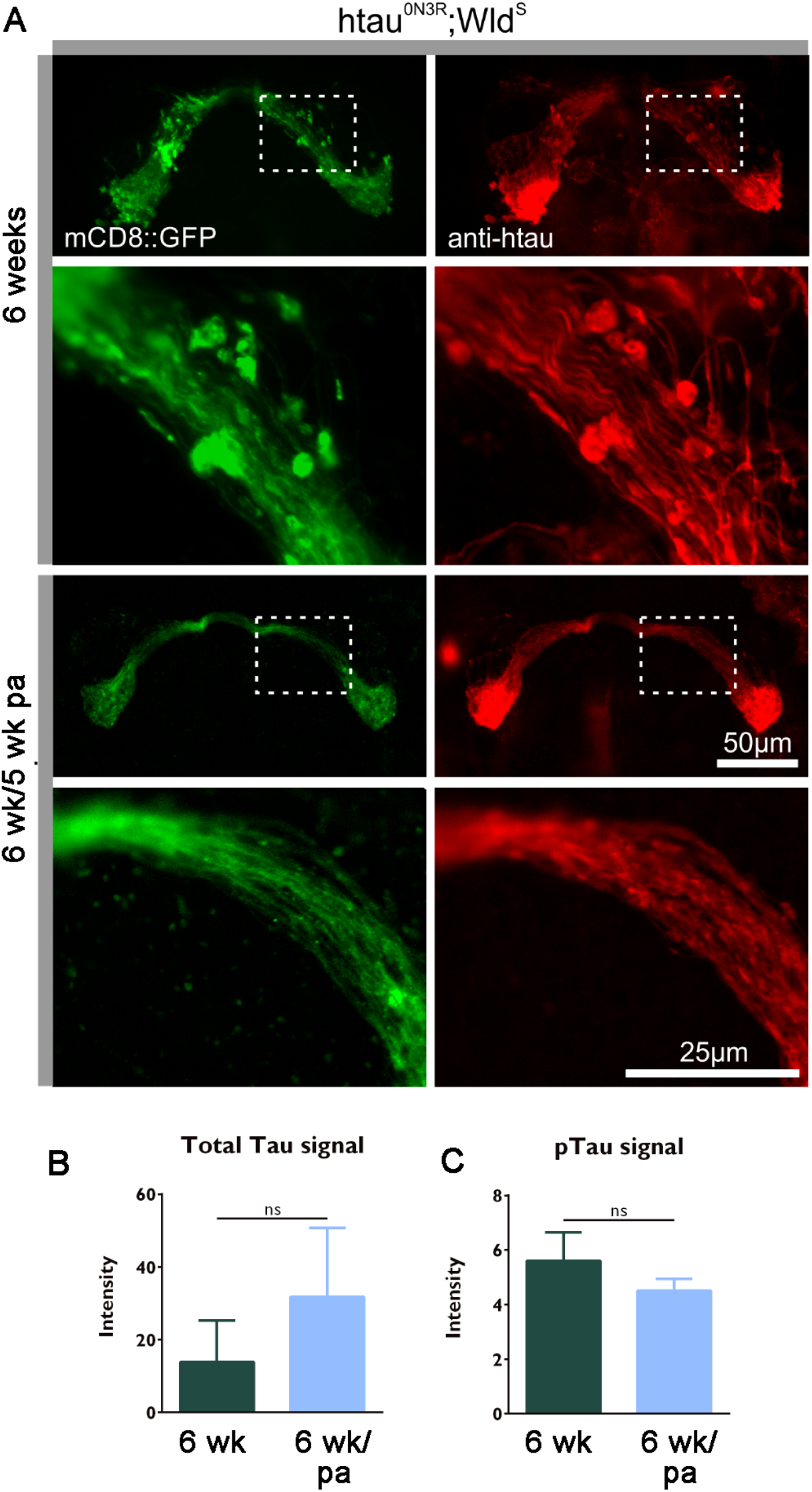
Tau expression in naïve hTau^0N3R^;Wld^S^ expressing axons is not different to hTau^0N3R^;Wld^S^ axons where Wld^S^ pathway has been activated. Visualisation of the membrane bound CD8-GFP protein shows extensive membrane fragmentation indicative of axonal degeneration in naïve hTau^0N3R^;Wld^S^ neurons at 6 weeks (green upper panels in A). Human tau is found within both the axonal processes as well as axonal swellings as visualised by a polyclonal anti-tau antibody (red upper panels in A). No such membrane fragmentation is evident in 6 week old hTau^0N3R^;Wld^S^ expressing axons (green lower panel in A) even 5 weeks after Wld^S^ pathway activation (pa) despite persisting human tau levels (red lower panels in A). (B) Quantification shows no differences in levels of total human tau though there is a (non-significant) trend for a reduction in tau phosphorylated at PHF-1 between naïve hTau^0N3R^;Wld^S^ expressing axons and hTau^0N3R^;Wld^S^ expressing axons where Wld^S^ has been activated even 6 weeks post activation (pa). (n=5). Values are presented as the mean ± SEM (p>0.05).

Since hyper-phosphorylation has been shown to mediate tau toxicity in many *Drosophila* models (5) (7) (44), it is conceivable that the activated Wld^S^-pathway is altering the degenerative changes by reducing the levels of phosphorylated hTau. To investigate this, hTau phosphorylated at the PHF-1 site was quantified in hTau^0N3R^;Wld^S^ axons with and without Wld^S^-pathway activation. There was a trend for a reduction in the PHF-1 signal in the hTau^0N3R^;Wld^S^ flies where Wld^S^ was activated but this was not significant (Fig 5c).

## Discussion

The axonal compartment of neurons is susceptible to tau-mediated dysfunction and degeneration making it a potential therapeutic target in the treatment of neurodegenerative disease. This study demonstrates that when the pathway downstream of Wld^S^ is “activated” in hTau^0N3R^;Wld^S^ axons, tau-mediated axonal swellings *were prevented from forming*. Significantly, in animals allowed to develop axonal swellings due to hTau^0N3R^ expression any further progression of pathology was halted after the Wld^S^-pathway was activated. This protective effect was seen without alterations in total tau and importantly was dependent on the activation of pathways downstream of Wld^S^. Understanding the mechanisms by which activation of the Wld^S^-protective pathway negates tau-mediated axonal degeneration could yield important insight into how axons degenerate in tauopathy and other similar chronic degenerative conditions, and provide novel disease-modifying targets that emulate this protective effect.

### Variable impact of Wld^S^ overexpression in previous models of neurodegeneration: can this be explained by the need to “activate” pathway downstream of Wld^S^ to uncover neuroprotection

Previous studies of Wld^S^ in models of chronic neurodegeneration have indicated that expression of Wld^S^ has variable effects on disease phenotypes. This was also evident in our study where no rescue of hTau^0N3R^-mediated neuronal dysfunction or degeneration was seen in either larvae or adults stages following mere co-expression of Wld^S^. Similar results were reported in the *SOD1-G93A* model of MND (45). Like this, there is a large body of conflicting evidence of Wld^S^ sensitivity in chronic neurodegenerative diseases displaying Wallerian-like degeneration. Wld^S^ did not alter axonal degeneration in models of prion disease (46), MND (45) (47) and hereditary spastic paraplegia (48). In contrast, in models that investigate degeneration with an acute onset, such as toxic neuropathy (28), ischaemic injury (49) and MPTP induced-Parkinsonism (24) (50) a protective effects of Wld^S^ is reported.

One interpretation of the dissimilar effect of Wld^S^ in models of acute and chronic neurodegeneration and their variable sensitivity to Wld^S^ could be a different mechanism of axonal degeneration occurring in acute compared with chronic neurodegenerative conditions. Another explanation for the variable sensitivity to Wld^S^ in models of chronic neurodegeneration could simply be that its protective effect is a general delaying of degeneration which is not always apparent in the time period assayed in the chronic models in question. Our data imply that this is unlikely to be the case since no protective effect emerged at even very late time points when Wld^S^ was simply expressed with hTau^0N3R^ (Fig 1). Instead, we propose and that another explanation may be provided by a key difference between the experimental paradigms employed to study acute degeneration, which is missing in the models of chronic neurodegeneration. This is that in all acute models, injury has to be simulated to create the acute condition and this may set in motion a series of events that “activate” the Wld^S^ protective pathway. This is never done in models of chronic neurodegeneration so it is conceivable that in those models the protective effect of Wld^S^ is not induced due to inadequate “activation” of the pathway that Wld^S^ acts upon. This would limit the impact of Wld^S^ on the ensuing neurodegeneration. Where there is partial rescue of phenotype in chronic models, the Wld^S^ pathway may start to become “activated” as the degeneration sets in. “Activation” of the Wld^S^ pathway by simulating injury or established neurodegeneration is a novel concept. There is no precedence for this idea from studies published to date because no one has reported overlaying an acute injury in a chronic model. Our data indicates that Wld^S^ behaves differently in uninjured axons compared to injured ones – the mechanisms responsible for this need to be elucidated.

### Activation of the Wld^S-^protective pathway prevents as well as halts progression of tau-mediated degeneration

The axonal degeneration observed in hTau^0N3R^ transgenic flies was characterised by axonal swellings, which are indicative of the early stages of axonal degeneration caused by human tau. Axonal swellings are not a feature of Wallerian degeneration in the peripheral nervous system, however they have been described following injury in the CNS (13) (51). Axonal swellings have also been observed in models of neurodegeneration, including AD (51) (52) (53) and tauopathy (54) (9), leading to suggestions that Wallerian-like degeneration is occurring in neurodegenerative diseases.

Upon activation of the Wld^S^ pathway, we did not observe axonal swellings and there was a lack of any other feature of tau-mediated degeneration in hTau^0N3R^;Wld^S^ axons. These tau-expressing axons looked as normal as the Wld^S^ expressing controls, which previous studies have shown to be both morphologically normal, as well as physiologically functional (30). This protective effect was evident even when the Wld^S^ pathway was activated in the hTau^0N3R^;Wld^S^ axons at a time point after tau-mediated degeneration was established. This demonstrates that once activated, the Wld^S^ protective pathway prevents emergence of tau-mediated degeneration as well as halting progression of already established degeneration.

### What is the mechanism by which activated Wld^S^ pathway protects against hTau?

Tau-mediated degeneration is dependent upon factors including total tau levels (55), phosphorylation at pathological sites (56) (35) and tau aggregation (57). The presence of human tau within hTau^0N3R^;Wld^S^ axons, even weeks after activation of the Wld^S^ pathway, indicates that the protection seen was not due to a reduction in total tau level as a result of loss of human tau from the axon. Nor is it likely to be due to any significant reduction in its phosphorylation status at the one pathological site, PHF1 (ser/thr 396 and 404) that we examined. This surprising observation implies that upon activation, the Wld^S^-pathway acts to negate the degenerative effects of tau, despite the persistence of pathologically phosphorylated human tau within the axon. Nonetheless it is possible that this protective effect was conferred by reduced misfolding or phosphorylation at other pathological sites that have previously been implicated in tau-mediated degeneration in other *Drosophila* models of tauopathy, such as MC1 (57), AT100 (Thr212/Ser214) (58) or 12E8 (ser262/356) (35) or reduced aggregation of human tau remains to be determined by future studies. It is conceivable that Wld^S^ pathway activation may influence tau phosphorylation at other sites as well as modulate tau aggregation, as several isoforms of NMNAT, one of which is a component of Wld^S^ fusion protein, have been shown to act as potent chaperones of phosphorylated tau, preventing its aggregation *in vitro* (37). This report, and others like it which implicate NMNAT in modulation of tau-mediated toxicity in other experimental models (38) (39), suggest that activation of Wld^S^ pathway does not act downstream of tau, and instead there are some key points of intersection.

A better understanding of the mechanisms that underpin the Wld^S^ protective pathway may highlight commonalities with hTau mediated degeneration and thus identify potential points of intersection at which protection is conferred. A key component of the Wld^S^ pathway is the NAD^+^ salvage pathway, which has recently also been linked with AD (59). Wld^S^ contains nicotinamide mononucleotide adenylyl transferase 1 (NMNAT1) (60) (61), which is the final enzyme in the NAD^+^ salvage pathway in mammals, and the biosynthetic activity of NMNAT1 is required for the full Wld^S^ protective phenotype (62) (63). One isoform in mammals (NMNAT2) and the sole *Drosophila* homolog dNMNAT are rapidly lost upon injury and this is associated with degeneration (64). Wld^S^ is believed to compensate for the loss of NMNAT in injured Wld^S^ expressing axons, thereby preventing loss of NAD^+^ and activation of the downstream pro-degenerative pathway. dSarm/Sarm1 (32) and Highwire/PHR1 (34) are endogenous mediators of axon degeneration that affect levels of NMNAT, with Axundead (30) and Pebbled (65) identified downstream of the loss of NMNAT. Knockout of these endogenous mediators results in axon survival after injury. We postulate that the Wld^S^ protective pathway would not be activated in uninjured hTau;Wld^S^ expressing axons, and so its protective effects would not be evident. Upon activation, the Wld^S^ pathway may confer protection against tau-mediated degeneration because its downstream components are switched on and can interact directly with pathological tau. Supporting this, it has been shown that NMNAT prevents tau-mediated aggregation *in vitro*, acts as chaperone for proteosis of tau in rodent models and promotes clearance of tau oligomers leading to suppression of tau-induced degeneration in *Drosophila* models of tauopathy (38) (37, 39) (39). Alternatively, protection may be conferred indirectly due to downstream neuroprotective effects of Wld^S^ pathway activation, such as enhanced mitochondrial calcium buffering (66), or reduced oxidative stress (67) negating tau-mediated dysregulation in intracellular calcium and/or tau-mediated mitochondrial dysfunction (68) (59) or oxidative stress (43). Future studies will be required to identify the exact point at which the activated Wld^S^ pathway intersects with and therefore protects against tau-mediated degeneration. In particular it will be vital to explore whether expression of downstream meditors of the Wld^S^ pathway that potentially block injury-induced axon degeneration (such as NMNAT, dSarm, axed, highwire, or even NAD^+^) also modulate tau-mediated degeneration to emulate Wld^S^ pathway activation.

## Conclusion

We show that activation of the Wld^S^-pathway reliably protects against tau-mediated axonal degeneration, almost abolishing it. It is vital to understand how engagement of this pathway and its downstream mediators interact with tau, whether directly or indirectly, to halt tau-mediated axonal degeneration. This will yield important clues about the mechanisms underpinning tau-mediated axonal degeneration as well as enable identification of novel drug targets that can emulate the complete protection we report here, to truly halt tau-mediated degeneration in all tauopathies.

## Acknowledgements

This study was funded by The Gerald Kerkut Charitable Trust.

